# Natural and Pathogenic Protein Sequence Variation Affecting Prion-Like Domains Within and Across Human Proteomes

**DOI:** 10.1101/626648

**Authors:** Sean M. Cascarina, Eric D. Ross

## Abstract

Protein aggregation is involved in a variety of muscular and neurodegenerative disorders. For many of these disorders, current models suggest a prion-like molecular mechanism of disease, whereby proteins aggregate and spread to neighboring cells in an infectious manner. A variety of proteins with prion-like domains (PrLDs) have recently been linked to these disorders. The development of prion prediction algorithms has facilitated the large-scale identification of PrLDs among “reference” proteomes for various organisms. However, the degree to which intraspecies protein sequence diversity influences predicted aggregation propensity for PrLDs has not been systematically examined. Here, we explore protein sequence variation introduced at genetic, post-transcriptional, and post-translational levels, and its influence on predicted aggregation propensity for human PrLDs. We find that sequence variation is relatively common among PrLDs and in some cases can result in relatively large differences in predicted aggregation propensity. Analysis of a database of sequence variants associated with human disease reveals a number of mutations within PrLDs that are predicted to increase aggregation propensity. Our analyses expand the list of candidate human PrLDs, estimate the effects of sequence variation on the aggregation propensity of PrLDs, and suggest the involvement of prion-like mechanisms in additional human diseases.

## INTRODUCTION

Prions are infectious proteinaceous elements, most often resulting from the formation of self-replicating protein aggregates. A key component of protein aggregate self-replication is the acquired ability of aggregates to catalyze the conversion of identical proteins to the non-native, aggregated form. Although prion phenomena may occur in a variety of organisms, budding yeast has been used extensively as a model organism to study the relationship between protein sequence and prion activity (1–4). Prion domains from yeast prion proteins tend to share a number of unusual compositional features, including high glutamine/asparagine (Q/N) content and few charged and hydrophobic residues (2, 3). Furthermore, the amino acid composition of these domains (rather than primary sequence) is the predominant feature conferring prion activity (5, 6). This observation has contributed to the development of a variety of composition-centric prion prediction algorithms designed to identify and score proteins based on sequence information alone (7–13).

Many of these prion prediction algorithms were extensively tested and validated in yeast as well. For example, multiple yeast proteins with experimentally-demonstrated prion activity were first identified as high-scoring prion candidates by early prion prediction algorithms (9–11). Synthetic prion domains, designed *in silico* using the <u>P</u>rion <u>A</u>ggregation <u>P</u>rediction <u>A</u>lgorithm (PAPA), exhibited *bona fide* prion activity in yeast (14). Additionally, application of these algorithms to proteome sequences for a variety of organisms has led to a number of important discoveries. The first native bacterial PrLDs with demonstrated prion activity in bacteria (albeit in an unrelated bacterial model organism) were also initially identified using leading prion prediction algorithms (15, 16). A prion prediction algorithm was used in the initial identification of a PrLD from the model plant organism *Arabidopsis thaliana* (17), and this PrLD was shown to aggregate and propagate as a prion in yeast (though it is currently unclear whether it would also have prion activity in its native host). Similarly, multiple prion prediction algorithms applied to the *Drosophila* proteome identified a prion-like domain with *bona fide* prion activity in yeast (18). A variety of PrLD candidates have been identified in eukaryotic virus proteomes using prion prediction algorithms (19), and one viral protein was recently reported to behave like a prion in eukaryotic cells (20). These examples represent vital advances in our understanding of protein features conferring prion activity, and illustrate the broad utility of prion prediction algorithms.

Some prion prediction algorithms may even have complementary strengths: identification of PrLD candidates with the first generation of the <u>P</u>rion-<u>L</u>ike <u>A</u>mino <u>A</u>cid <u>C</u>omposition (PLAAC) algorithm led to the discovery of new prions (11), while application of PAPA to this set of candidate PrLDs markedly improved the discrimination between domains with and without prion activity *in vivo* (7, 14). Similarly, PLAAC identifies a number of PrLDs within the human proteome, and aggregation of these proteins is associated with an assortment of muscular and neurological disorders (21–34). In some cases, increases in aggregation propensity due to single amino acid substitutions are accurately predicted by multiple aggregation prediction algorithms, including PAPA (33, 35). Furthermore, the effects of a broad range of mutations within PrLDs expressed in yeast can also be accurately predicted by PAPA and other prion prediction algorithms, and these predictions generally extend to multicellular eukaryotes, albeit with some exceptions (36, 37). The complementary strengths of PLAAC and PAPA are likely derived from their methods of development. The PLAAC algorithm identifies PrLD candidates by compositional similarity to domains with known prion activity, but penalizes all deviations in composition (compared to the training set) regardless of whether these deviations enhance or diminish prion activity. PAPA was developed by randomly mutagenizing a canonical Q/N-rich yeast prion protein (Sup35) and directly assaying the frequency of prion formation, which was used to quantitatively estimate of the prion propensity of each of the 20 canonical amino acids. Therefore, PLAAC seems to be effective at successfully identifying PrLD candidates, while PAPA is ideally-suited to predict which PrLD candidates are most likely to have true prion activity, and how changes in PrLD sequence might affect prion activity.

To date, most proteome-scale efforts of prion prediction algorithms have focused on the identification of PrLDs within reference proteomes (i.e. a representative set of protein sequences for each organism). However, reference proteomes do not capture the depth and richness of protein sequence variation that may affect PrLDs *within* a species. Here, we explore the depth of intraspecies protein sequence variation affecting human PrLDs at the genetic, post-transcriptional, and post-translational levels. We estimate the range of aggregation propensity scores resulting from known protein sequence variation, for all high scoring PrLDs. To our surprise, aggregation propensity ranges are remarkably large, suggesting that natural sequence variation could potentially result in large inter-individual differences in aggregation propensity for certain proteins. Furthermore, we define a number of proteins whose aggregation propensities are affected by alternative splicing or pathogenic mutation. In addition to proteins previously linked to prion-like disorders, we identify a number of high-scoring PrLD candidates whose predicted aggregation propensity increases for certain isoforms or upon mutation, and some of these candidates are associated with prion-like behavior *in vivo* yet are not currently classified as “prion-like”. Finally, we provide comprehensive maps of PTMs within human PrLDs derived from a recently-collated PTM database.

## RESULTS

### Sequence Variation in Human PrLDs Leads to Wide Ranges in Estimated Aggregation Propensity

Multiple prion prediction algorithms have been applied to specific reference proteomes to identify human PrLDs (13, 38-42). While these predictions provide important baseline maps of PrLDs in human proteins, they do not account for the considerable diversity in protein sequences across individuals. In addition to the ~42k unique protein isoforms (spanning ~20k protein-encoding genes) represented in standard human reference proteomes, the human proteome provided by the neXtProt database includes >6 million annotated single amino acid variants (43). Importantly, these variants reflect the diversity of human proteins, and allow for the exploration of additional sequence space accessible to human proteins.

The majority of known variants in human coding sequences are rare, occurring only once in a dataset of ~60,700 human exomes (44). However, the frequency of multiple-variant co-occurrence for each possible variant combination in a single individual has not been quantified on a large scale. Theoretically, the frequency of rare variants would result in each pairwise combination of rare variants occurring in a single individual only a few times in the current human population. We emphasize that this is only a rough estimate, as it assumes independence in the frequency of each variant, and that the observed frequency of rare variants corresponds to the actual population frequency.

With these caveats in mind, we applied a modified version of our <u>P</u>rion <u>A</u>ggregation <u>P</u>rediction <u>A</u>lgorithm (PAPA; see Methods for modifications and rationale) to the human proteome reference sequences to obtain baseline aggregation propensity scores and to identify relatively high-scoring PrLD candidates. Since sequence variants could increase predicted aggregation propensity, we employed a conservative aggregation propensity threshold (PAPA score≥0.0) to define high-scoring PrLD candidates (*n* = 5173 unique isoforms). Nearly all PrLD candidates (*n* = 5028; 97.2%) have at least one amino acid variant that influenced the PAPA score, indicating that the variant mapped specifically to the PrLD region. Protein sequences for all pairwise combinations of known protein sequence variants were computationally generated for all proteins with moderately high-scoring PrLDs (>20million variant sequences, derived from the 5173 protein isoforms with PAPA score≥0.0). While most proteins had relatively few variants that influenced predicted aggregation propensity scores, a number of proteins had >1000 unique PAPA scores, indicating that PrLDs can be remarkably diverse (Fig 1A). To estimate the overall magnitude of the effects of PrLD sequence variation, the PAPA score range was calculated for each set of variants (i.e. for all variants corresponding to a single protein). PAPA score ranges adopt a right-skewed distribution, with a median PAPA score range of 0.10 (Fig 1B,C; Table S1). Importantly, the estimated PAPA score range for a number of proteins exceeds 0.2, indicating that sequence variation can have a dramatic effect on predicted aggregation propensity (by comparison, the PAPA score range=0.92 for the entire human proteome). Additionally, we examined the aggregation propensity ranges of prototypical prion-like proteins associated with human disease (21-25, 27-34), which are identified as high-scoring candidates by both PAPA and PLAAC. In most cases, the lowest aggregation propensity estimate derived from sequence variant sampling scored well-below the classical aggregation threshold (PAPA score=0.05), and the highest aggregation propensity estimate scored well-above the aggregation threshold (Fig 1D). Furthermore, for a subset of prion-like proteins (FUS and hnRNPA1), aggregation propensity scores derived from the initial reference sequences differed considerably for alternative isoforms of the same protein, suggesting that alternative splicing may also influence aggregation propensity. It is possible that natural genetic variation between individuals may substantially influence the prion-like behavior of human proteins.

**Fig 1.**
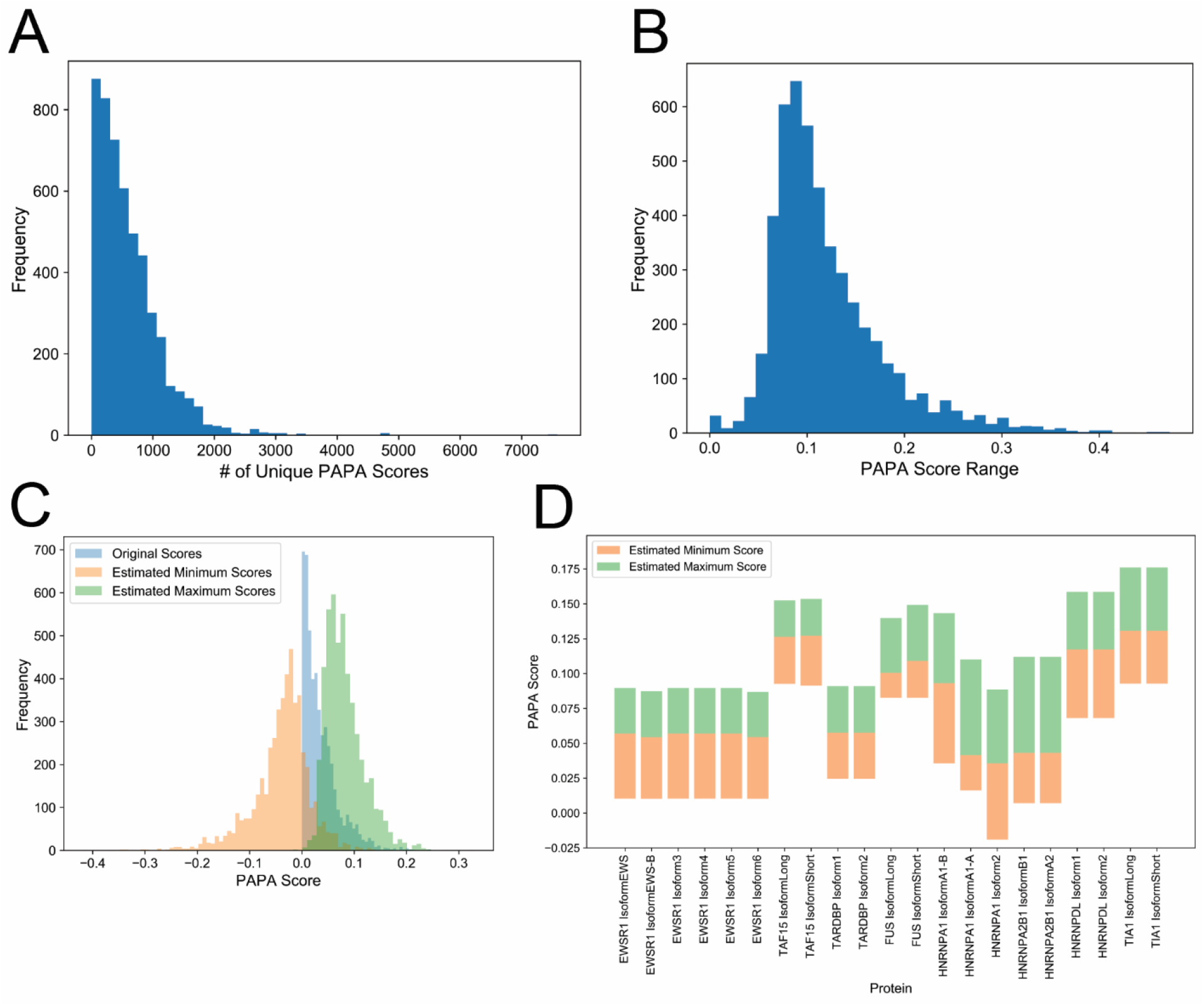
Sampling of human PrLD sequence variants yields broad ranges of aggregation propensity scores. (A) Histogram indicating the frequencies corresponding to the number of unique PAPA scores per protein. (B) The distribution of aggregation propensity ranges, defined as the difference between the maximum and minimum aggregation propensity scores from sampled sequence variants, is indicated for all PrLDs scoring above PAPA=0.0 and with at least one annotated sequence variant. (C) Histograms indicating categorical distributions of aggregation propensity scores for the theoretical minimum and maximum aggregation propensity scores attained from PrLD sequence variant sampling, as well as original aggregation propensity scores derived from the corresponding reference sequences. (D) Modified box plots depict the theoretical minimum and maximum PAPA scores (lower and upper bounds, respectively), along with the reference sequence score (the color transition point) for all isoforms of prototypical prion-like proteins associated with human disease.

### Alternative Splicing Introduces Sequence Variation that Affects Human PrLDs

As observed in Fig 1D, protein isoforms derived from the same gene can correspond to markedly different aggregation propensity scores. Alternative splicing essentially represents a form of post-transcriptional sequence variation *within* each individual. Alternative splicing could affect aggregation propensity in two main ways. First, alternative splicing could lead to the inclusion or exclusion of an entire PrLD, which could modulate prion-like activity in a tissue-specific manner, or in response to stimuli affecting the regulation of splicing. Second, splice junctions that bridge short, high-scoring regions could generate a complete PrLD, even if the short regions in isolation are not sufficiently prion-like.

The ActiveDriver database (45) is a centralized resource containing downloadable and computationally accessible information regarding “high-confidence” protein isoforms, post-translational modification sites, and disease associated mutations in human proteins. We first examined whether alternative splicing would affect predicted aggregation propensity for isoforms that map to a common gene. In total, of the 39532 high-confidence isoform sequences, 8018 isoforms differ from the highest-scoring isoform mapping to the same gene (Table S2). Most proteins maintain a low aggregation propensity score even for the highest-scoring isoform. However, we found 159 unique proteins for which both low-scoring and high-scoring isoforms exist (Fig 2A; 414 total isoforms that differ from the highest-scoring isoform), suggesting that alternative splicing could affect prion-like activity. Furthermore, it is possible that known, high-scoring prion-like proteins are also affected by alternative splicing. Indeed, 15 unique proteins had at least one isoform that exceeded the PAPA threshold, and at least one isoform that scored even higher (Fig 2B). Therefore, alternative splicing may affect aggregation propensity for proteins that are already considered high-scoring PrLD candidates.

**Fig 2.**
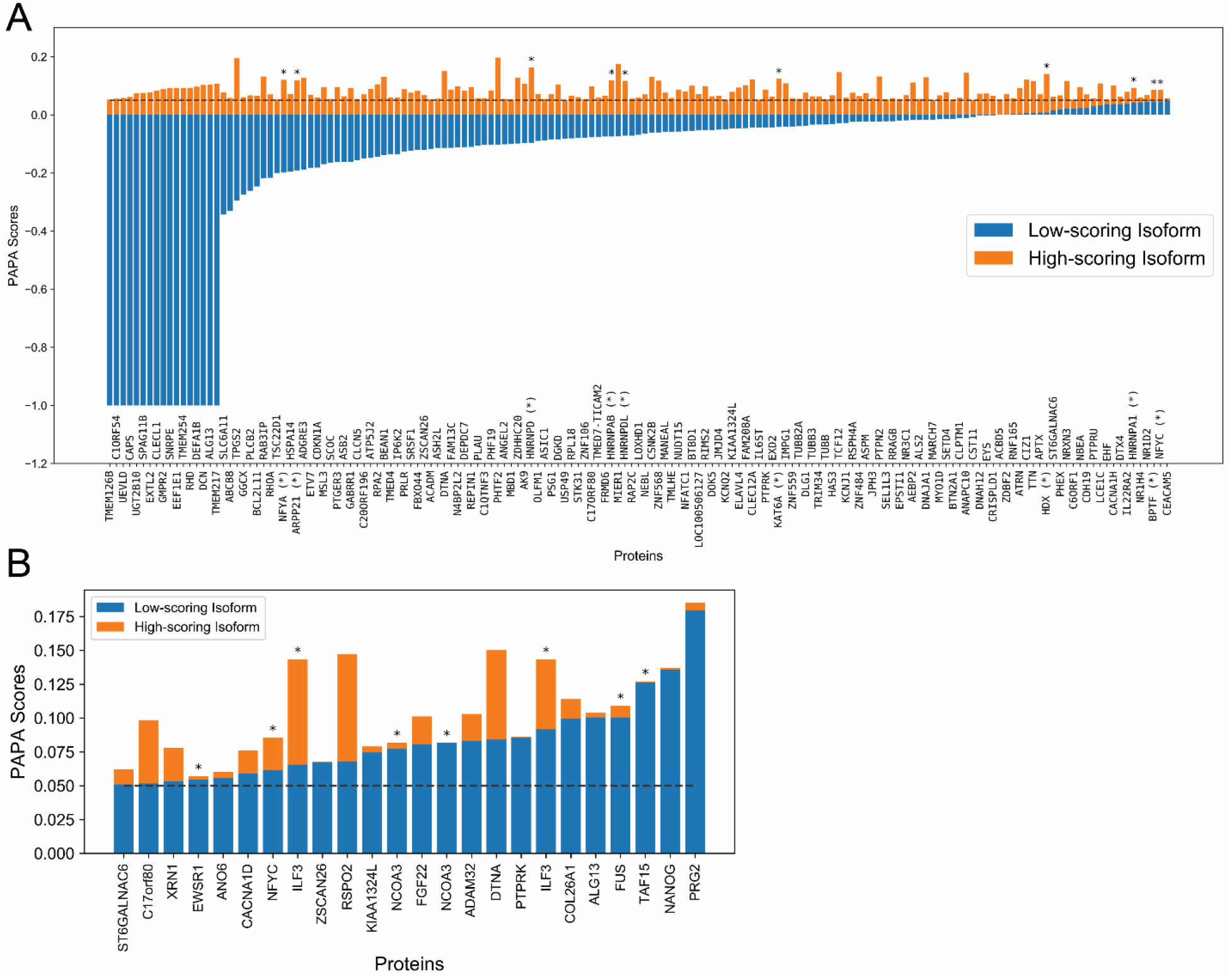
Alternative splicing influences predicted aggregation propensity for a number of human PrLDs. (A) Minimum and maximum aggregation propensity scores (indicated in blue and orange respectively) are indicated for all proteins with at least one isoform below the classical PAPA=0.05 threshold and at least one isoform above the PAPA=0.05 threshold. For simplicity, only the highest and lowest PAPA score are indicated for each unique protein (*n*=159), though many of the indicated proteins that cross the 0.05 threshold have multiple isoforms within the corresponding aggregation propensity range (*n*=414 total isoforms; Table S2). (B) For all protein isoforms with an aggregation propensity score exceeding the PAPA=0.05 threshold and with at least one higher-scoring isoform (*n*=48 total isoforms, corresponding to 15 unique proteins), scores corresponding to the lower-scoring and higher-scoring isoforms are indicated in blue and orange respectively. In both panels, asterisks (*) indicate proteins for which a PrLD is also identified by PLAAC. Only isoforms for which splicing affected the PAPA score are depicted.

Strikingly, many of the prototypical disease-associated prion-like proteins were among the high-scoring proteins affected by splicing. For example, hnRNPDL, which is linked to limb girdle muscular dystrophy type1G, has one isoform scoring far below the 0.05 PAPA threshold and another scoring far above the 0.05 threshold. hnRNPA1, which is linked to a rare form of myopathy and to amyotrophic lateral sclerosis (ALS), also has one isoform scoring below the 0.05 PAPA threshold and one isoform scoring above the threshold. Additionally, multiple proteins linked to ALS, including EWSR1, FUS, and TAF15 all score above the 0.05 PAPA threshold and have at least one isoform that scores even higher. Mutations in these proteins are associated with neurological disorders involving protein aggregation or prion-like activity. Therefore, in addition to well-characterized mutations affecting aggregation propensity of these proteins, alternative splicing may play an important and pervasive role in disease pathology, either by disrupting the intracellular balance between aggregation-prone and non-aggregation-prone variants, or by acting synergistically with mutations to further enhance aggregation propensity.

The fact that numerous proteins already linked to prion-like disorders have PAPA scores affected by alternative splicing raises the intriguing possibility that additional candidate proteins identified here may be involved in prion-like aggregation under certain conditions or when splicing is disrupted. For example, the RNA-binding protein XRN1 is a component of processing-bodies (or “P-bodies”), and can also form distinct synaptic protein aggregates known as “XRN1 bodies”. Prion-like domains have recently been linked to the formation of membraneless organelles, including stress granules and P-bodies (46). Furthermore, dysregulation of RNA metabolism, mRNA splicing, and the formation and dynamics membraneless organelles are prominent features of prion-like disorders (46). However, XRN1 possesses multiple low-complexity domains that are predicted to be disordered, so it will be important to determine which (if any) of these domains are involved in prion-like activity. Interestingly, multiple β-tubulin proteins (TUBB, TUBB2A, and TUBB3) are among proteins with both low-scoring and high-scoring isoforms. Expression of certain β-tubulins is misregulated in some forms of ALS (47, 48), β-tubulins aggregate in mouse models of ALS (49), mutations in α-tubulin subunits can directly cause ALS (50), and microtubule dynamics are globally disrupted in the majority of ALS patients (51). The nuclear transcription factor Y subunits NFYA and NFYC, which both contain high-scoring PrLDs affected by splicing, are sequestered in Htt aggregates in patients with Huntington’s disease (52). NFYA has also been observed in aggregates formed by the TATA-box binding protein, which contains a polyglutamine expansion in patients with spinocerebellar ataxia 17 (53). HNRNP A/B constitutes a specific member of the hnRNP A/B family, and encodes both a low-scoring and a high-scoring isoform. The high-scoring isoforms resembles prototypical prion-like proteins, containing two RNA-recognition motifs (RRMs) and a C-terminal PrLD (which is absent in the low-scoring isoform, and hnRNP A/B proteins were shown to co-aggregate with PABPN1 in a mammalian cell model of oculopharyngeal muscular dystrophy (54). Alternative splicing of ILF3 mRNA leads to the direct inclusion or exclusion of a PrLD in the resulting protein isoforms NFAR2 and NFAR1, respectively (55, 56). NFAR2 (but not NFAR1) is recruited to stress granules, its recruitment is dependent upon its PrLD, and recruitment of NFAR2 leads to stress granule enlargement (57). A short “amyloid core” from the high-scoring NFAR2 PrLD forms amyloid fibers *in vitro* (41). ILF3 proteins co-aggregate with mutant p53 (another PrLD-containing protein) in models of ovarian cancer (58). ILF3 proteins are also involved in the inhibition of viral replication upon infection by dsRNA viruses, re-localize to the cytoplasm in response to dsRNA transfection (simulating dsRNA viral infection), and appear to form cytoplasmic inclusions (59). Similarly, another RNA-binding protein, ARPP21, is expressed in two isoforms: a short isoform containing two RNA-binding motifs (but lacking a PrLD), and a longer isoform containing both RNA-binding motifs as well as a PrLD. The longer isoform (but not the short isoform) is recruited to stress granules, suggesting that the recruitment is largely dependent on the C-terminal PrLD (60). Furthermore, most of the proteins highlighted above have PrLDs that are detected by both PAPA and PLAAC (Table S2), indicating that these results are not unique to PAPA.

Collectively, these observations suggest that alternative splicing may play an important and pervasive role in regulating the aggregation propensity of certain proteins, and that misregulation of splicing could lead to an improper intracellular balance of a variety of aggregation-prone isoforms.

### Disease-Associated Mutations Influence Predicted Aggregation Propensity for a Variety of Human PrLDs

Single-amino acid substitutions in prion-like proteins have already been associated with a variety of neurological disorders (46). However, the role of prion-like aggregation/progression in many disorders is a relatively recent discovery, and additional prion-like proteins continue to emerge as key players in disease pathology. Therefore, the list of known prion-like proteins associated with disease is likely incomplete, and raises the possibility that PrLD-driven aggregation influences additional diseases in currently undiscovered or underappreciated ways.

We leveraged the ClinVar database of annotated disease-associated mutations in humans to examine the extent to which clinically-relevant mutations influence predicted aggregation propensity within PrLDs. For simplicity, we focused on single-amino acid substitutions that influenced aggregation propensity scores. Of the 33059 single-amino acid substitutions (excluding mutation to a stop codon), 2385 mutations increased predicted aggregation propensity (Table S3). Of these proteins, 27 unique proteins scored above the 0.05 PAPA threshold and had mutations that increased predicted aggregation propensity (83 total mutants), suggesting that these mutations lie within prion-prone domains and are suspected to enhance protein aggregation (Fig 3A). Additionally, 24 unique proteins (37 total mutants) scored below the 0.05 PAPA threshold but crossed the threshold upon mutation (Fig 3B).

**Fig 3.**
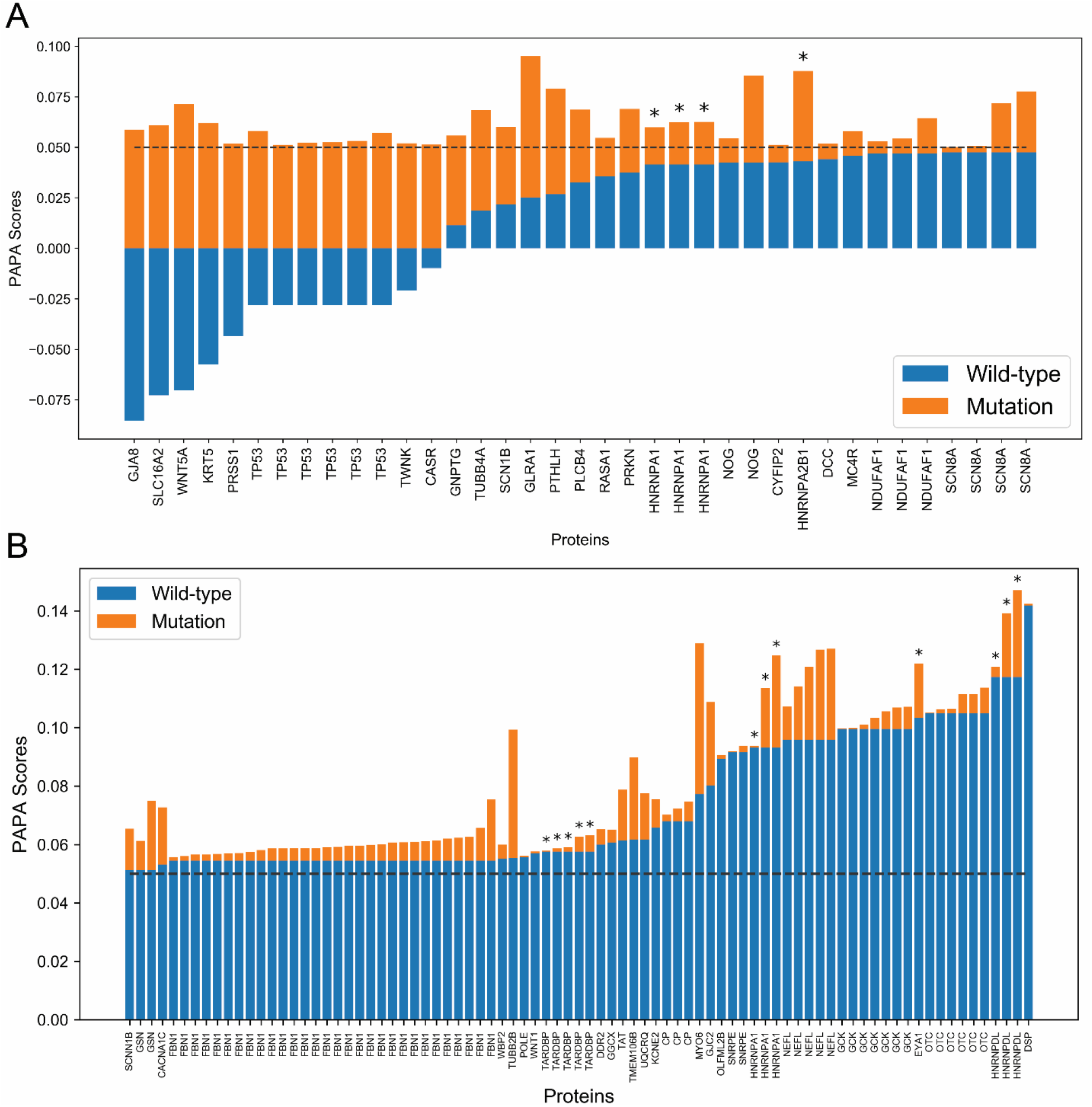
Disease-associated mutations influence predicted aggregation propensities of known PrLDs and new candidate prion-like proteins. (A) For all disease-associated single-amino acid substitutions that map to high-scoring PrLDs (PAPA score>0.05) and increase predicted aggregation propensity score, scores corresponding to the wild-type and mutant sequences are indicated in blue and orange respectively. (B) Wild-type and mutant aggregation propensity scores are similarly plotted for all proteins with wild-type PAPA score <0.05 but mutant PAPA score >0.05. In both panels, asterisks (*) indicate proteins also containing a PLAAC-positive PrLD.

As observed for protein isoforms affecting predicted aggregation propensity, a number of mutations affecting prion-like domains with established roles in protein aggregation associated with human disease (21–25, 27–34, 61) were among these small subsets of proteins, including TDP43, hnRNPA1, hnRNPDL, hnRNPA2B1, and p53. However, a number of mutations were also associated with disease phenotypes that have not currently linked to prion-like aggregation. For example, in addition to hnRNPA1 mutations linked to prion-like disorders (which are also detected in our analysis; Fig3, and Table S3), K277N, P275S, and P299L mutations in the hnRNPA1 PrLD increase its predicted aggregation propensity yet are associated with chronic progressive multiple sclerosis (Table S3), which is currently not considered a prion-like disorder. It is possible that, in addition to known prion-like disorders, certain forms of progressive multiple sclerosis (MS) may also involve prion-like aggregation. Intriguingly, the hnRNPA1 PrLD (which overlaps with its M9 nuclear localization signal) is targeted by autoantibodies in MS patients (62), and hnRNPA1 mislocalizes to the cytoplasm and aggregates in patients with MS (63), similar to observations in hnRNPA1-linked prion-like disorders (33).

Many of the high scoring proteins with mutations affecting aggregation propensity have been linked to protein aggregation, yet are not currently considered prion-like. For example, missense mutations in the PrLD of light chain neurofilament protein (encoded by the NEFL gene) are associated with autosomal dominant forms of Charcot-Marie Tooth (CMT) disease (64). Multiple mutations within the PrLD are predicted to increase aggregation propensity (Fig 3A and Table S3), and a subset of these mutations have been shown to induce aggregation of both mutant and wild-type neurofilament light protein in a dominant manner in mammalian cells (65). Fibrillin 1 (encoded by the FBN1 gene) is a structural protein of the extracellular matrix that forms fibrillar aggregates as part of its normal function. Mutations in fibrillin 1 are predominantly associated with Marfan Syndrome, and lead to connective tissue abnormalities and cardiovascular complications (66). While the majority of disease-associated mutations affect key cysteine residues (Table S3), a subset of mutations lie within its PrLD and are predicted to increase aggregation propensity (Fig 3A), which could influence normal aggregation kinetics, thermodynamics, or structure. Multiple mutations within the PrLD of the gelsolin protein (derived from the GSN gene) are associated with Finnish type familial amyloidosis [also referred to as Meretoja syndrome; (67–69)] and are predicted to increase aggregation propensity (Fig 3A). Furthermore, mutant gelsolin protein is aberrantly proteolytically cleaved, releasing protein fragments that overlap with the PrLD and are found in amyloid deposits in affected individuals [for review, see (70)].

For proteins that cross the classical 0.05 aggregation propensity threshold, proteins exhibiting large relative changes in predicted aggregation propensity upon single-amino acid substitution likely reflect changes in intrinsic disorder classification implemented in PAPA via the FoldIndex algorithm. Therefore, these substitutions may reflect the disruption of predicted structural regions, thereby exposing high-scoring PrLD regions normally buried in the native protein. Indeed multiple mutations in the prion-like protein p53 lead to large changes in predicted aggregation propensity (Fig 3B, Table S3), are thought to disrupt p53 structural stability, and result in a PrLD that encompasses multiple predicted aggregation-prone segments (71). Additionally, two mutations in the Parkin protein (encoded by the PRKN/PARK2 gene), which has been linked to Parkinson’s disease, increase its predicted aggregation propensity (Fig 3B, Table S3). Parkin is prone to misfolding and aggregation upon mutation (72, 73) and in response to stress (74, 75). Indeed, both mutants associated with an increase in predicted aggregation propensity for Parkin were shown to decrease Parkin solubility, and one of the mutants forms microscopically-visible foci in mammalian cells (72). It is important to note that, while both mutations that increase predicted aggregation propensity disrupt the catalytic site of Parkin, aggregation of Parkin may also contribute to disease pathology.

### A Survey of Post-translational Modifications Within Human PrLDs

Post-translational modifications (PTMs) represent a form of protein sequence variation in which the intrinsic properties of amino acids in synthesized proteins are altered via chemical modification. Recently, information derived from multiple centralized PTM resources, as well as individual studies, have been combined into a single database describing a broad range of PTM sites across the human proteome (45). PTMs could directly affect protein aggregation by increasing or decreasing inherent aggregation propensity. Indeed, changes in PTMs have been associated with a variety of aggregated proteins in neurodegenerative diseases (76–78), and PTMs can influence liquid-liquid phase separation (79), which has recently been linked to low-complexity domains and PrLDs. Therefore, PTMs likely play an important role in regulating the aggregation propensity of certain PrLDs.

Using centralized PTM databases, we mapped PTMs to human PrLDs. While the contribution of each of the canonical amino acids to aggregation of PrLDs has been fairly well-characterized (7, 80), consistent effects of each type of PTM on aggregation of PrLDs have not been defined. Therefore, we mapped PTMs to PrLDs using a relaxed aggregation propensity threshold (PAPA cutoff=0.0, rather than the standard 0.05 threshold), which accounts for the possibility that PTMs could increase aggregation propensity or regulate the solubility of proteins whose aggregation propensity is near the standard 0.05 aggregation threshold.

For each PTM type, distributions for the number of modifications per PrLD are shown in Fig 4A, and PTMs mapped to PrLDs are provided in Table S4. Although PTMs are likely important regulators of aggregation for certain PrLDs and should be examined experimentally on a case-by-case basis, we explored whether any PTMs were globally enriched or depleted within PrLDs. Since PrLDs typically have unusual amino acid compositions (which would affect the gross total for some PTMs within PrLDs), the number of potentially modifiable residues for each type of PTM was first calculated for the whole proteome and for PrLDs and statistically compared (see Methods for detailed description).

**Fig 4.**
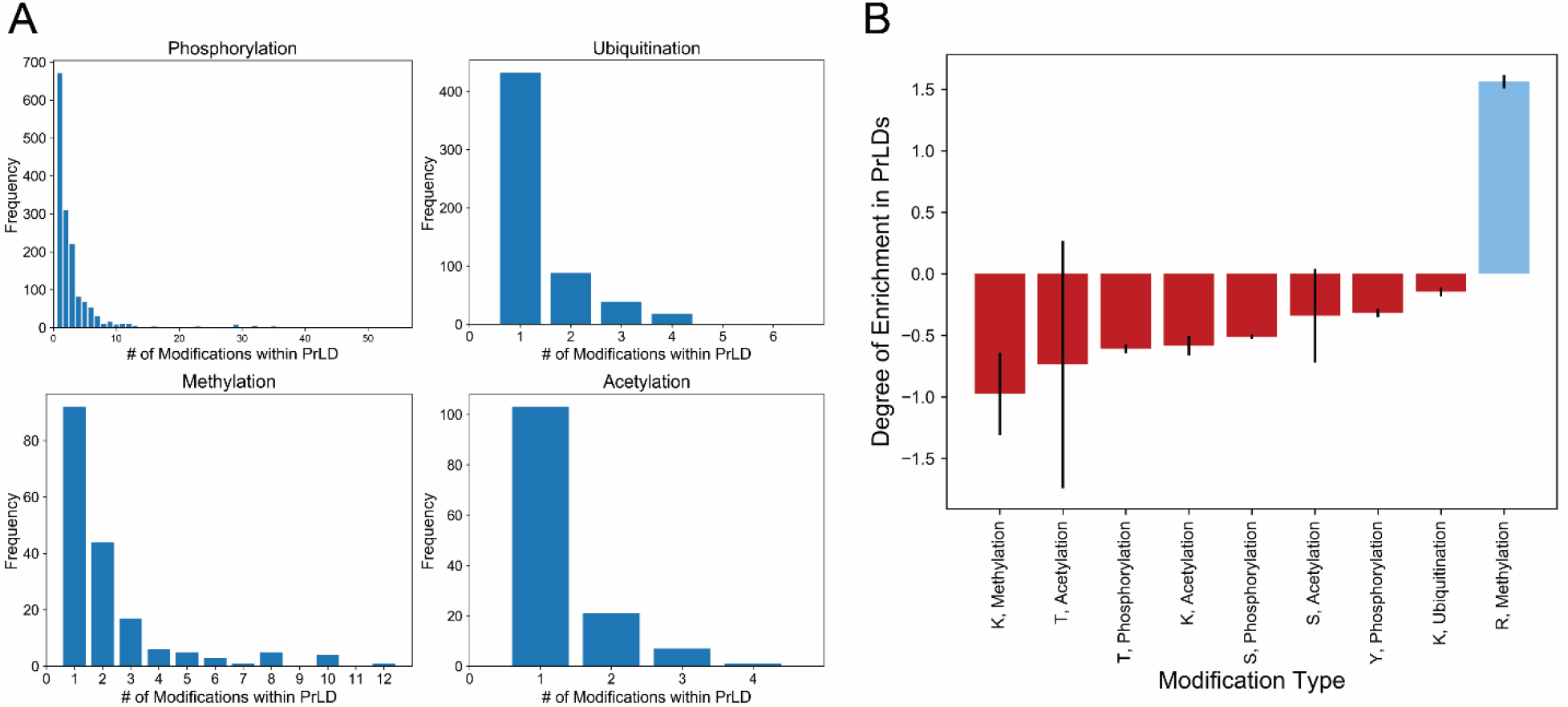
Certain PTM types are enriched or depleted within human PrLDs. (A) Distributions depicting the number of modifications within each PrLD for each of the main PTM types. (B) Estimated degree of enrichment (blue) or depletion (red) for each PTM type within human PrLDs. Error bars represent the standard error.

Arginine methylation was the only PTM type significantly enriched in human PrLDs (Fig 4B and Table S5). In contrast, serine phosphorylation, threonine phosphorylation, tyrosine phosphorylation, lysine acetylation, lysine methylation, and lysine ubiquitination are significantly depleted within human PrLDs. The global underrepresentation of nearly all PTM types within PrLDs is particularly surprising since PrLDs are typically intrinsically disordered, and many of the PTM types studied here are enriched within intrinsically disordered regions *vis-à-vis* ordered regions (81). However, this may be reconciled by considering the amino acid compositions associated with the flanking regions surrounding PTM sites. For example, regions flanking phosphorylation sites are typically enriched in charged residues and depleted in neutral and aromatic residues (82). Similarly, the flanking regions of arginine methylation sites are significantly associated with increased net charge and high glycine content (among other properties) and decreased glutamine and glutamic acid content (83). Regions flanking lysine methylation sites are also enriched in glycine, aromatic residues, and threonine, and depleted in non-aromatic hydrophobic residues, glutamine, and glutamic acid. This highlights an important point: while these features are consistent with PTM sites occurring preferentially within intrinsically disordered regions, they may be specific for disordered regions of particular amino acid compositions. Therefore, although PrLDs are typically considered intrinsically disordered, the Q/N-richness of most PrLDs may result in fewer PTMs compared to non-Q/N-rich intrinsically disordered regions.

Nevertheless, the global depletion of PTMs within PrLDs does not imply a lack of importance for PTMs that do occur within PrLDs. The mapping of PTMs to PrLDs may catalyze the experimental determination of the effects of each individual PTM on PrLD aggregation.

### Sequence Variation at the Genetic, Transcriptional, and Posttranslational Levels is Associated with Disease-Relevant Aggregation of a PrLD-Containing Protein – A Case Study of hnRNPA1

We were surprised to find that the hnRNPA1 PrLD is affected by every form of sequence variation examined in the present study, including genetic variation, alternative splicing, multiple disease-associated mutations, and post-translational modification (Fig 5). The short isoform, hnRNPA1-A (320 amino acids), scores just below the 0.05 PAPA threshold. Multiple mutations within the hnRNPA1 PrLD increase prion propensity and *in vivo* aggregation (33). The long isoform, hnRNPA1-B (372 amino acids), scores substantially higher than the short isoform (PAPA scores are 0.093 and 0.042, respectively), and contains the region affected by the disease-associated mutations. It is possible that mutations within the hnRNPA1 PrLD, in combination with the high scoring isoform, have particularly potent aggregation-promoting effects. Under the current model for prion-like aggregation, the high-scoring protein isoform [which is typically less-abundant than the low-scoring isoform (84, 85)] could “seed” protein aggregates, which may then be capable of recruiting the lower-scoring isoform. Although this is currently speculative, it is supported by a recent study, which showed that mutation in the TDP-43 PrLD and cytoplasmic aggregation of TDP-43 in ALS patients was associated with dysregulation of hnRNPA1 mRNA splicing (85, 86). This dysregulation led to increased abundance of the high-scoring hnRNPA1-B isoform and subsequent aggregation of the hnRNPA1 protein (85). Finally, 31 unique posttranslational modifications map to the hnRNPA1 long-isoform PrLD, particularly to sites immediately flanking the highest-scoring PrLD region. It may also be possible that perturbations in posttranslational regulation of hnRNPA1, could influence protein aggregation *in vivo*. For example, phosphorylation of certain modification sites within the hnRNPA1 PrLD are differentially modified upon osmotic shock, which promotes accumulation of hnRNPA1 in the cytoplasm (87), and a variety of PTMs within the PrLD regulate additional aspects of hnRNPA1 localization and molecular interactions (88). Together, these observations suggest that multiple types of sequence variation may conspire to simultaneously influence hnRNPA1-related disease phenotypes.

**Fig 5.**
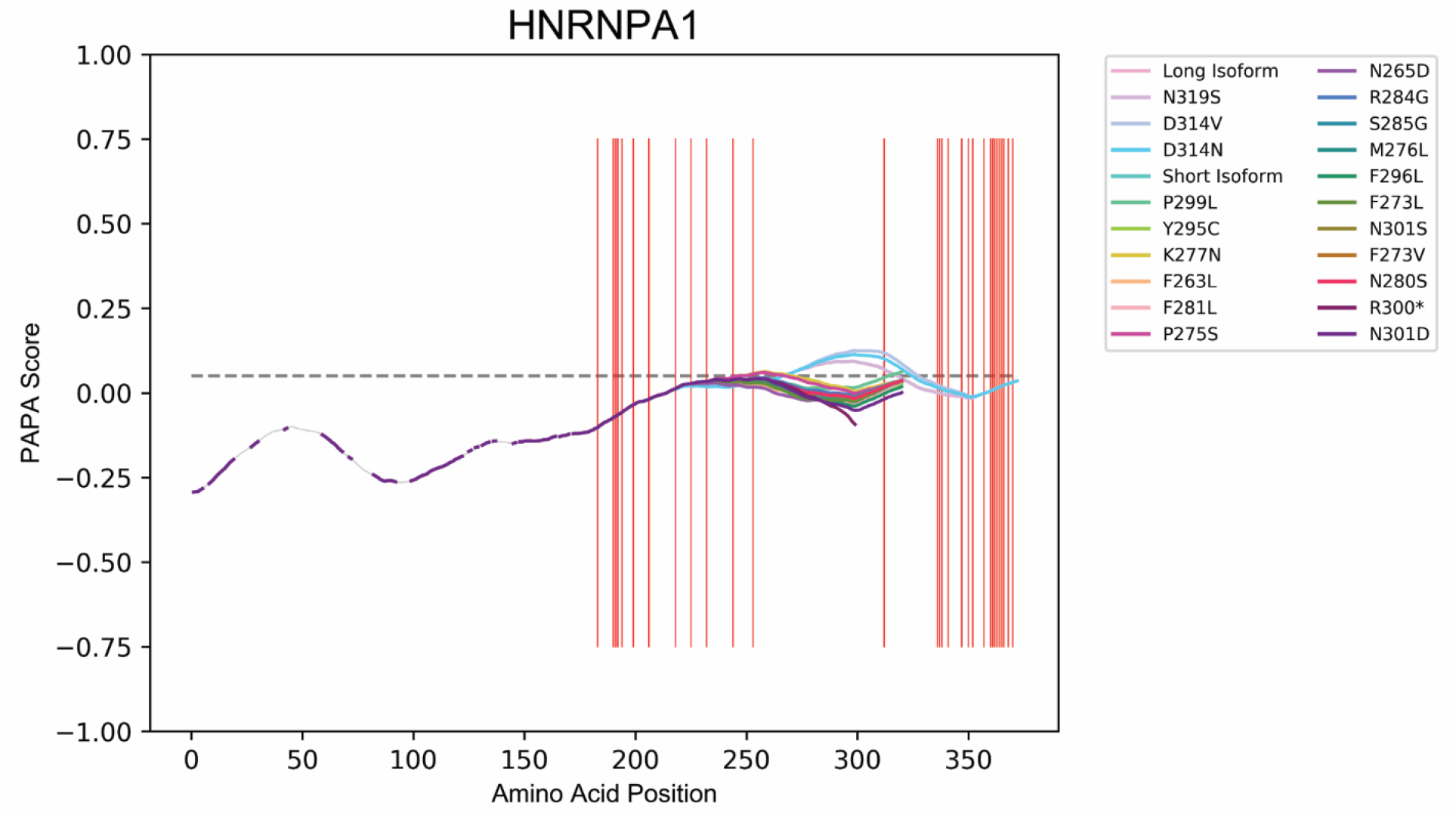
The hnRNPA1 PrLD is affected by genetic, post-transcriptional, and post-translational sequence variation. Aggregation propensity scores for all hnRNPA1 splice variants, as well as all disease-associated variants, are plotted separately. For each line, regions corresponding to FoldIndex scores >0.0 (which are not assigned aggregation propensity scores in PAPA) are plotted as thin grey segments, whereas all regions scored by PAPA (FoldIndex<0.0) are plotted as thick colored segments. All PTMs mapping to regions with relatively high-scoring regions (PAPA>0.0) are indicated by vertical red lines. The classical PAPA=0.05 threshold is indicated with a dashed grey line.

## DISCUSSION

Numerous studies have explored the pervasiveness of candidate PrLDs across a variety of organisms. Although initial prediction of prion propensity among reference proteomes is an important first step in identifying candidate PrLDs, these predictions do not account for the richness of sequence diversity across individuals of the same species. Here, we complement these studies with an in-depth analysis of human intraspecies sequence variation and its effects on predicted aggregation propensity for PrLDs.

Prion aggregation is strongly (though not exclusively) dependent on the physicochemical characteristics of the aggregating proteins themselves. While analyses of reference proteomes necessarily treat protein sequences as invariable, protein sequence variation can be introduced at the gene, transcript, or protein levels via mutation, alternative splicing, or post-translational modification, respectively. Importantly, these protein changes can exert biologically-relevant effects on protein structure, function, localization, and physical characteristics, which could influence prion-like behavior.

Broadly, we found that protein sequence variation is common within human PrLDs, and can influence predicted aggregation propensity rather substantially. Using the frequency of observed single-amino acid variants from a large collection of human exomes (~60,700 individuals), we estimated the range of aggregation propensity scores by generating all pairwise combinations of variants for moderately high-scoring proteins. Aggregation propensity score ranges were often remarkably large, indicating that sequence variation could, in theory, have a dramatic effect on the prion-like behavior of certain proteins. However, it is important to note that not all variant combinations may naturally occur. For example, it is possible that certain variants commonly co-occur *in vivo*, or that some variants are mutually exclusive. Indeed, it is likely that aggregation propensity acts as a selective constraint which limits the allowable sequence space that can be viably explored by PrLDs. Conversely, our method conservatively assumed that all single amino acid variants were rare, even though some variants are substantially more common (44): it is possible that some double, triple, or even quadruple variants may occur in a single individual with some regularity. Therefore, while our method for sampling sequence variants may over- or under-estimate aggregation propensity ranges for some PrLDs, our results nevertheless highlight the sequence diversity within PrLD regions across individuals. In principle, subtle changes in prion-like behavior could have phenotypic consequences, and may explain at least a small portion of human phenotypic diversity, although we emphasize that this is currently speculative.

We also identified a variety of proteins for which alternative splicing influences predicted aggregation propensity, which has a number of important implications. According to the prion model of protein aggregation, it is possible that aggregation of high-scoring isoforms could seed the aggregation of lower-scoring isoforms, assuming at least a portion of the PrLD is present in both isoforms. Importantly, this “cross-seeding” could occur even if the aggregation propensity of the low-scoring isoform is not itself sufficient to promote aggregation. Additionally, tissue-specific expression or splicing of certain proteins could impact prion-like behavior, effectively compartmentalizing or modulating prion-like activity in specific tissues. This also implies that dysregulation of alternative splicing could lead to overproduction of aggregation-prone isoforms. Interestingly, many of the prion-like proteins found in aggregates in individuals with neurological disease are splicing factors, and their sequestration into aggregates may impact the splicing of mRNAs encoding other aggregation-prone proteins (85). This was recently proposed to produce a “snowball effect”, whereby aggregation of key proteins result in the aggregation of many other proteins via an effect on splicing or expression which could, in-turn, affect the aggregation of additional proteins (89).

Protein sequence variation can be beneficial, functionally inconsequential, or pathogenic. Examination of pathogenic sequence variants specifically (i.e. mutations in PrLDs associated with human disease) yielded a number of new prion-like protein candidates. Many of these new candidates have been associated with protein aggregation in previous studies, yet are not widely classified as prion-like, making them perhaps the most promising candidates for future studies and in-depth experimentation. In addition to candidates with experimental support, a number of candidates have not been previously linked to prion-like activity but may still have yet undiscovered prion-like activity *in vivo*. It is worth noting that, while PAPA and PLAAC predictions often overlap, many of these new candidate PrLDs (when considering disease-associated mutations) were only identified by PAPA, so experimental confirmation of aggregation and prion-like behavior is necessary.

Post-translational modification represents the final stage at which cells can modify protein properties and behavior. In a number of cases, PTMs are associated with protein aggregation across a diverse set of neurodegenerative disorders (76–78). However, the precise effects of PTMs on aggregation propensity and whether they play a causative role in protein aggregation are often unclear. Nevertheless, one could speculate about what the effects of each PTM might be with respect to aggregation of PrLDs based on prion propensities for the 20 canonical amino acids and the physicochemical characteristics of the PTM. For example, charged residues typically inhibit prion aggregation within PrLDs (7, 80), so phosphorylation of serine, threonine, or tyrosine residues may tend to suppress prion-like activity (90). Conversely, lysine acetylation or N-terminal acetylation neutralizes the charge, increases hydrophobicity, and introduces hydrogen bond acceptors, which may positively contribute to prion activity. Arginine and lysine methylation does not neutralize the charge, but slightly increases the bulkiness and hydrophobicity of the sidechain. Asymmetric dimethylation of arginine is common within proteins with PrLDs (91) and can weaken cation-pi interactions with aromatic sidechains within PrLDs (92). Recent studies implicate arginine methylation (which was the only PTM type significantly enriched within human PrLDs in our study) as an important suppressor of PrLD phase separation and pathological aggregation [for review, see (79, 91)]; together with our data, this suggests that arginine methylation may play a vital role in regulating the aggregation propensity of a multitude of PrLDs. Ubiquitination of lysine residues within PrLDs may sterically hinder PrLD aggregation. There are likely additional considerations that extend beyond the physicochemical properties of PTMs that alter aggregation propensity. For example, the proportion of any particular PrLD-containing protein that is modified at a given time in the cell dictates the effective concentration of each species which may influence the likelihood of forming a stable aggregate, analogous to the apparent resistance to prion disease in humans that are heterozygous at position 129 in the prion protein, PrP (93). PTMs also regulate subcellular localization, protein-protein interactions, and structural characteristics, which may secondarily influence PrLD aggregation propensity. As with any attempt at generalizing predictions, the effects of PTMs may be highly context-specific, depending on interactions with particular neighboring residues. To facilitate further exploration of PTMs within PrLDs, we mapped PTMs from collated PTM databases to human PrLDs, and provide these maps as resources to encourage case-by-case experimental exploration.

As a final note, we would like to emphasize caution in over-interpreting our observations. As mentioned above, prion-like activity *in vivo* is strongly dependent upon the physicochemical characteristics of PrLDs, which are largely determined by protein sequence. However, prion-like aggregation can be influenced *in vivo* by factors other than inherent sequence characteristics, including expression levels, subcellular localization, protein chaperone activity, and molecular binding partners, among others (94). The influence of these factors will likely vary on a case-by-case basis; two similarly aggregation-prone PrLDs may be differentially regulated, leading one to aggregate while the other remains functional/soluble. At the same time, our prion prediction algorithm was developed in the context of a eukaryotic model organism (7), thereby incorporating at least some contribution from additional cellular factors and a crowded intracellular environment. Furthermore, prion-like aggregation is one of many possible mechanisms that can affect protein function upon mutation or alternative splicing. We are not advocating for a mutual exclusivity view of prion-like aggregation: protein sequence variation can have multiple concomitant consequences, and prion-like aggregation may simply be one of those consequences. For example, mutations can disrupt native protein sequence, resulting in loss of function of the protein. But those same mutations may also enhance prion-like aggregation, leading to a cytotoxic gain-of-function and a contribution to overall disease pathology. Additionally, while we have focused in this study on mutations that increase predicted aggregation propensity, mutations within PrLDs that decrease predicted aggregation propensity may be equally important. Adaptive, reversible aggregation activity exhibited by some PrLDs may involve a delicate balance in kinetic and thermodynamic parameters, which could be disrupted by mutations that either decrease or increase predicted prion-like behavior. Mutations that decrease predicted aggregation propensity may ultimately lead to PrLD aggregation *in vivo* if the loss in inherent aggregation propensity is ultimately outweighed by an indirect increase in aggregation propensity caused, for example, by disrupted molecular interactions that normally sequester the PrLD. Therefore, sequence variants that affect high-scoring PrLDs yet decrease predicted aggregation propensity may still be of interest and utility, and are retained in all supplementary resources.

Collectively, our results shed new light on the relationship between protein sequence diversity and inherent aggregation propensity, highlight a number of promising new prion-like candidates whose aggregation propensities may be influenced by protein sequence variation, and provide a variety of resources to propel future protein aggregation research.

## METHODS

### Data Acquisition and Processing

Human protein isoform sequences, along with corresponding clinical variants [originally derived from NCBI’s ClinVar database (95, 96)] and PTM sites, were acquired from the ActiveDriver database [(45); https://www.activedriverdb.org/; downloaded on 10/5/2018]. For estimation of the range of theoretical aggregation propensity scores based on observed sequence variants, reference sequences including >6 million annotated single amino acid variants were obtained from the neXtProt database [(43, 97); https://www.nextprot.org/; downloaded on 2/12/2019].

All data processing, including data re-structuring, quantification, calculation, statistical analysis, and plotting was performed using in-house Python scripts. All statistical analyses were performed using the built-in Python stats module with default settings, except that all statistical tests were two-sided. Where applicable, correction for multiple hypothesis testing was implemented via the statsmodels package available for Python. All plotting was performed using the Matplotlib and Seaborn packages.

### Modifications to the Original PAPA Algorithm

PAPA source code was downloaded (http://combi.cs.colostate.edu/supplements/papa/) and augmented with custom functions scripted in Python. Briefly, the original PAPA algorithm assigns aggregation propensity scores to each position in a protein based on a combined score from 41 consecutive 41-amino acid windows (effectively, an 81-amino acid window for each position) (7, 98). Our modified PAPA algorithms differs from the original PAPA algorithm in three key ways: 1) PAPA scores are assigned to the last residue of the first sliding window, which improves the scoring of protein termini and is critical for mapping PTM sites to PrLDs; 2) overlapping domains within a single protein that exceed a pre-defined PAPA threshold are merged, which yields precise definitions of predicted PrLD boundaries and accounts for multiple PrLDs within a single protein; and 3) predictions of protein disorder are simplified by calculating the FoldIndex over each full window, rather than the average of 41 consecutive windows. Additionally, for many analyses, a relaxed aggregation propensity threshold of 0.0 was chosen for two main reasons: 1) sequence variation or post-translational modification may increase aggregation propensity in some cases, such that the aggregation propensity may lie beyond our classical 0.05 threshold upon modification or mutation, and 2) this threshold captures ~10% of each proteome, yielding a reasonable set of high-scoring proteins for analysis.

### Estimation of Aggregation Propensity Ranges Via Exhaustive Pairwise Variant Combination

All possible pairwise combinations of single amino acid variants (neXtProt database) within proteins with a relatively high baseline aggregation propensity (PAPA score>0.0) were generated computationally and stored as independent sequences. Theoretical sequence variants were then scored using our modified PAPA algorithm, and the minimum, maximum, and reference sequence scores were subsequently compared. By default, PAPA assigns an arbitrary score of −1.0 to proteins lacking a predicted intrinsically disordered region. Therefore, variants with a theoretical minimum PAPA score of −1.0 were excluded from analyses.

### Analysis of PTM Enrichment/Depletion within PrLDs

PrLDs are, by definition, biased in terms of amino acid composition (2, 3). Without controlling for compositional biases, certain PTMs would be over- or under-represented among PrLDs simply by virtue of the availability of modifiable residues. Therefore, when comparing protein modifications within PrLDs vs. the remainder of the proteome, non-modified residues were defined as residues capable of being modified by the PTM of interest but with no empirical evidence of modification. For example, serine phosphorylation was analyzed by comparing the number of phosphorylated serine residues within PrLDs to the number of non-phosphorylated serine residues within PrLDs. Calculations were performed similarly for non-PrLD regions (i.e. the remainder of the proteome). The degree of PTM enrichment within PrLDs was then calculated as:

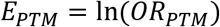

and

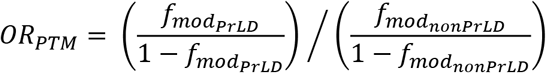

where 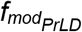 and 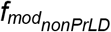 represent the fraction of modified residues out of potentially modifiable residues for the given PTM type within PrLD and non-PrLD regions, respectively. Statistical enrichment or depletion for each PTM type within PrLDs was evaluated using a two-sided Fisher’s exact test, with Benjamini-Hochberg correction for multiple hypothesis testing (with false discovery rate threshold of 0.05).

## DATA ACCESS

All data generated or analyzed during this study are included in this published article as supplementary information, or the original sources have been described and cited.

## ACKNOWLEDGEMENTS

We would like to thank members of the Ross lab for helpful discussion and insight.

## Supplementary Table Legends

**Table S1. Minimum, maximum, and reference sequence PAPA scores derived from random sampling of sequence variant combinations for proteins with high-scoring PrLDs.** For all proteins with a moderately high-scoring PrLD (PAPA>0.0) and at least one single-amino acid variant, the minimum and maximum aggregation propensity scores obtained from randomly sampling PrLD sequence variants, along with the corresponding reference sequence score, are indicated (see Methods for a complete description of sequence variant calculations).

**Table S2. Aggregation propensity scores and inter-isoform score comparison for all human protein isoforms.** Predicted aggregation propensity for all “high-confidence” human protein isoforms (derived from ActiveDriverDB) was calculated using the modified PAPA algorithm (see Methods section for details). Scores and corresponding full protein sequences are indicated for all isoforms, along with the maximum PAPA score among all isoforms mapping to the same gene, the difference between the PAPA score for the indicated isoform and the maximum PAPA score among related isoforms, and the protein sequence corresponding to the highest-scoring related isoform. Additionally, the PLAAC algorithm was used to analyze the same sequences. A binary variable indicates if the protein contains a PLAAC-predicted PrLD that overlaps with the PAPA-predicted PrLD for high-scoring proteins only and, if so, the position of the PLAAC-predicted PrLD.

**Table S3. Comparison of wild-type and disease-associated mutant PAPA scores.** For all disease associated mutants in the ClinVar database, mutant sequences were generated by incorporating the indicated amino acid substitution at the appropriate position and re-scored using the modified PAPA algorithm. For each variant, both wild-type and mutant aggregation propensity scores are indicated (along with their corresponding protein sequences), as well as the difference between mutant and wild-type scores. For each variant, the associated disease phenotype annotation is also included. PLAAC predictions are also included, as indicated for Table S2.

**Table S4. Comprehensive mapping of PTMs within moderately high-scoring human PrLDs.** Human PTMs derived from the ActiveDriverDB were mapped to all human PrLDs with PAPA score>0.0. For each protein the maximum PAPA score, moderately high-scoring PrLD sequence (corresponding to all overlapping regions with PAPA score>0.0), amino acid positions bounding the PrLD sequence, and all PTMs mapping to the PrLD region are indicated. PLAAC predictions are also included, as indicated for Table S2.

**Table S5. Statistical analysis of PTM enrichment/depletion within human PrLDs.** For each PTM type, statistical enrichment or depletion within PrLDs was evaluated using a two-sided Fisher exact test (see Methods section for detailed description).

